# Age-related increases in PDE11A4 protein expression trigger liquid:liquid phase separation (LLPS) of the enzyme that can be reversed by PDE11A4 small molecules inhibitors

**DOI:** 10.1101/2024.10.01.616004

**Authors:** E Amurrio, J Patel, M Danaher, M Goodwin, P Kargbo, S Lin, CS Hoffman, S Ul Mahmood, DP Rotella, MP Kelly

## Abstract

PDE11A is a little-studied phosphodiesterase sub-family that breaks down cAMP/cGMP, with the PDE11A4 isoform being enriched in the memory-related brain region called the hippocampus. Age-related increases in PDE11A expression occur in human and rodent hippocampus and cause age-related cognitive decline of social memories. Interestingly, the age-related increase in PDE11A4 protein ectopically accumulates in spherical clusters that group together in the brain to form linear filamentous patterns termed “ghost axons.” The biophysical/physiochemical mechanisms underlying this age-related clustering of PDE11A4 are not yet known. As such, we determine here if age-related clustering of PDE11A4 may reflect liquid:liquid phase separation (LLPS), and if PDE11A inhibitors being developed for age-related cognitive decline can reverse this biomolecular condensation. We found that human and mouse PDE11A4 exhibit several LLPS-promoting sequence features including intrinsically disordered regions, non-covalent pi-pi interactions, and prion-like domains, with multiple bioinformatic tools predicting PDE11A4 undergoes LLPS. Consistent with these predictions, age-related PDE11A4 clusters were non-membrane bound spherical droplets that progressively fuse over time in a concentration-dependent manner. 5 different PDE11 inhibitors (tadalafil, BC11-38, SMQ-02-57, SMQ-03-30 and SMQ-03-20) across 3 scaffolds reversed PDE11A4 LLPS (a.k.a. remixing) in hippocampal HT22 cells, with PDE11A4 droplets reforming (a.k.a. de-mixing) following a 5-hour washout of low but not high concentrations of these compounds. Strikingly, a single oral administration of 30 mg/kg SMQ-03-20 substantially reduced the presence of PDE11A4 ghost axon in the aged mouse brain. Thus, PDE11A4 exhibits 4 defining criteria of LLPS, and PDE11A small molecule inhibitors reverse this age-related phenotype both *in vitro* and *in vivo*.

## INTRODUCTION

After the age of 60, nearly all individuals experience some form of cognitive decline—particularly memory deficits—and no drugs prevent or reverse this loss ^3, 4^. Indeed, advanced age is the strongest risk factor for dementia (e.g., ^10^). Even in absence of dementia, age-related cognitive impairment increases health care costs and risk for disability ^11^. Literature suggests intracellular cAMP and cGMP signaling are decreased in the aged and demented human and rodent hippocampus^12–14^. Age-related increases in phosphodiesterase 11A (PDE11A), an enzyme that breaks down cAMP/cGMP, are thought to contribute to these hippocampal signaling deficits and appear to be a fundamental mechanism underlying select long-term memory deficits associated with age-related cognitive decline^1, 15, 16^. That is, preventing age-related increases in PDE11A expression from occurring in mice prevents age-related cognitive decline of social memories ^1^, and reversing age-related increases in PDE11A4 protein in old mice reverses their memory deficits ^15^. Hence, we recently developed the first potent, selective and orally-bioavailable small molecule inhibitors of PDE11A for the treatment of age-related hippocampal pathologies^17, 18^.

PDE catalytic activity does not simply control the total cellular content of cyclic nucleotides, it generates individual subcellular signaling compartments^19, 20^. Such subcellular compartmentalization of cyclic nucleotides allows a single cell to respond discretely to diverse intra- and extracellular signals ^21^. Thus, the aberrant localization of a PDE has the potential to be far more damaging than a simple loss of catalytic activity^21^. Such mislocalization would not only remove a PDE from its normal pool(s) of cyclic nucleotides (i.e., a loss-of-function), it would potentially displace another PDE(s) and ectopically hydrolyze a foreign pool of cyclic nucleotides (i.e. a gain-of-function since different PDEs have differing catalytic activities). It is then quite interesting that age-related increases in PDE11A4 protein expression do not occur uniformly throughout the hippocampus, but rather are ectopically enriched in the ventral hippocampus (a.k.a., anterior hippocampus in primates) membrane fractions and “ghost axons”—that is, axonal terminals that are so densely packed with PDE11A4 protein that other axonal markers are occluded^1^.

Little is understood of the biophysical/physiochemical mechanisms regulating PDE11A4 age-related clustering, but liquid:liquid phase separation (LLPS) presents itself as a likely candidate. LLPS (a.k.a. biomolecular condensation or inclusion body formation) represents a reversible de-mixing event implicated in neurodegenerative disorders and aging^22^. Depending on the molecule, LLPS can sequester unneeded protein, buffer proteins (i.e., temporarily store and then release upon demand), or accelerate biochemical reactions by virtue of concentrating enzymes with substrates in membraneless organelles^23^. LLPS is driven by either heterotypic condensation (i.e., interaction between client and a scaffolding protein/RNA) or homotypic condensation (i.e., self-association), the latter of which is particularly well nucleated by intrinsically disordered regions (IDRs) ^22, 24^. Interestingly, both hPDE11A4 (https://www.ncbi.nlm.nih.gov/protein/NP_058649.3 accessed 08/15/24) and mPDE11A4 (https://www.ncbi.nlm.nih.gov/protein/NP_001074502.1 accessed 08/15/24) are annotated as containing an IDR within their regulatory N-terminal domain. Interestingly, phosphorylation of serines 117 and 124 (pS117/pS124) within this IDR promote age-related clustering of PDE11A4^1^ and phosphorylation of residues within IDRs are known to regulate LLPS^24, 25^. These physiochemical properties, in addition to the fact that PDE11A4 clusters are 1) largely spherical, 2) increase as endogenous expression increases, 3) require PDE11A4 homodimerization, and 4) fail to colocalize with typical organelle markers either *in vitro* or *in vivo* ^1, 15, 26^, strongly suggests that age-related clustering of PDE11A4 is caused by the protein undergoing LLPS. If so, it may be important to identify therapeutic approaches capable of reversing this LLPS. Thus, we sought to determine if 1) PDE11A4 does undergoes LLPS, and 2) if our PDE11A4-selective catalytic inhibitors are capable of reversing age-related clustering of PDE11A4 protein *in vitro* and *in vivo*. Here, we report that age-related clustering of PDE11A4 meets the criterion for LLPS, and that this LLPS is reversed by PDE11i’s across multiple scaffolds both *in vitro* and *in vivo*.

## METHODS

### LLPS Prediction

To determine if 1) PDE11A4 is likely to have additional IDRs outside of the N-terminal and 2) if any IDRs exhibit LLPS-promoting features we used the following reproducibly recommended^22, 27, 28^ computational screening tools: D2P2, https://d2p2.pro/search (accessed 8/29/23); PScore, http://abragam.med.utoronto.ca/~JFKlab/Software/psp.htm (accessed 8/29/23); PLAAC, http://plaac.wi.mit.edu (accessed 8/29/23); CIDER, https://pappulab.wustl.edu/CIDER/analysis (accessed 8/29/23; PhaSePred, http://predict.phasep.pro (accessed 8/31/23); and Phase Separation Predictor; http://www.pkumdl.cn:8000/PSPredictor (accessed 08/01/24).

### Plasmid generation

Plasmids were generated as previously described ^26^. Briefly, Genscript (Piscataway, NJ) generated constructs expressing either EmGFP alone containing an A206Y mutation to prevent EmGFP dimerization ^29^ or the mouse *Pde11a* (NM_001081033) sequence fused at the N-terminal with EmGFP. Note that mouse *Pde11a* is ∼95% homologous and the same length as human *Pde11a4* and so the protein is referred to herein as mPDE11A4 for clarity. These constructs were initially generated on a pUC57 backbone and then subcloned into a pcDNA3.1+ mammalian expression vector (Life Technologies; Waltham, MA).

### Cell culture and transfection

As described above, PDE11A4 expression in brain is enriched in the hippocampus ^16, 30, 31^. Therefore, we use the HT22 hippocampal cell line (sex undefined) for most *in vitro* investigations, with COS1 cells used solely for live-imaging given their larger size since they exhibit similar mPDE11A4 phenotypes as HT22 cells^1^. As previously described^1, 17^, cells were maintained in T-75 flasks in Dulbecco’s Modified Eagle Medium (DMEM) with sodium pyruvate (GIBCO, Gaithersburg, MD or Corning, Manassas, VA), 1% Penicillin/Streptomycin (P/S); GE Healthcare Life Sciences; Logan, UT), and 10% fetal bovine serum (FBS; Atlanta Biologicals), with incubators set to 37°C/5% CO_2_. Cells were passaged at ∼70% confluency using TrypLE Express (GIBCO; Gaithersburg, MD). The day before transfection, cells in DMEM+FBS+P/S were plated in 60 mm dishes, 24-well plates, or coverslips depending on the experiment. The day of transfection, the media was replaced with Optimem (GIBCO) and cells were transfected using 5 microliters Lipofectamine 2000 (Invitrogen; Carlsbad, CA) and 1.875 ug of plasmid DNA per 5 mL of optimem as per the manufacturer’s protocol. ∼19 hours post-transfection (PT), the Optimem/Lipofectamine solution was replaced with DMEM+FBS+P/S. Cells continued growing for five hours in the supplemented media and then followed 1 of 3 courses. They were either 1) stained with 2μM BODIPY-568 in (Invitrogen, #D3835) 1x phosphate-buffered saline (PBS 10x Powder Concentrate, Fisher BioReagents) for 15 minutes and then fixed in 4% paraformaldehyde and mounted using DAPI Fluoromount-G mounting media (Southern Biotech), 2) directly fixed in 4% paraformaldehyde (Sigma-Aldrich) in 1x PBS and then stored in 1x PBS, 2) pharmacologically treated for 1 hour or 24 hours and then fixed as above and stored in 1X PBS, or 3) treated for 1 hour or 24 hours, then switched to compound-free DMEM+FBS+P/S for 5 hours and then fixed and stored in 1X PBS. Images of cells labeled with Bodipy-568 were captured using the Leica Model DM6 CFS confocal microscope in the University of Maryland School of Medicine (UMSOM) Department of Neurobiology Imaging Core equipped with the Leica TCS SP8 laser system, Leica STP8000 control panel, Lumencor Sola Light Engine epifluorescent lamp, and Leica CTR6 power source using a 63X/1.40 Oil CS2 ∞/0.17/OFN25/E HC PL APO objective. Live cell imaging was conducted in DMEM+FBS+P/S at 37°C/5% CO_2_ on a Zeiss Axiovert 200M microscope in the University of South Carolina School of Medicine Instrumentation Resource Core Facility (RRID:SCR_024955). The Zeiss was equipped with a VivaTome Optical Train, an MRm AxioCam, a PeCon CTI Controller 3700 and a Fluar 20X/0.75 ∞/0.17 objective. Images were acquired every thirty seconds over the span of 45-120 minutes. Static images for quantification of mPDE11A4 compartmentalization into spherical clusters were collected from cells stored in 1x PBS using a Nikon Eclipse TE2000-E Inverted via a 10x/0.40 CS2 ∞/0.17/OFN25/A objective equipped with Photometrics CoolSNAP cf camera and CoolLED pE-300lite LED illuminator. Representative images for each well were captured using MetaVue v6.2r6 software and saved as jpeg files. This Nikon microscope is located in the UMSOM Department of Neurobiology Imaging Core. Over the course of experiments, cells were sporadically tested for yeast, fungal, and bacterial infections (Invitrogen; Cat#:C7028), with negative results always obtained.

### Compounds

Tadalafil (Tocris, 6311), BC11-38 (medchem express, Hy108618), rolipram (Sigma, R6520), and papaverine (Sigma, P3510) were obtained from commercial vendors. SMQ-02-57, SMQ-03-30, and SMQ-03-20 were synthesized by the Rotella lab as previously described^17, 18^. SMQ-02-57 is highly selective for PDE11A4 relative to other PDE families^17^. At 500 nM, SMQ-02-57 shows less than 20% inhibition of PDEs 3, 4, 5, 6 and 10. At 1 uM, it shows less than 15% inhibition of PDEs 1A, 2A, 7B, 8A and 9A^17^. SMQ-03-20 is also highly selective, showing ∼50% inhibition of PDE6C at 500 nM and 40% inhibition of PDE2A at 1 uM ^18^. SMQ-03-30 profiles similarly to SMQ-03-20 against these panels, with the addition of ∼50% inhibition of PDE10A at 1 uM ^18^.

### PDE activity assay

As previously described ^17, 18^, cells were treated for 1 hour, after which the media was removed, the cells were harvested in PDE assay buffer (20mM Tris-HCL and 10mM MgCl_2_) and homogenized using a tissue sonicator (output control: 7.5, duty cycle: 70, continuous). Samples were then held at 4 °C until processing. **T**otal protein levels were quantified using the DC Protein Assay Kit (Bio-Rad, Hercules, CA) according to the manufacturer’s directions. 3 μg of each sample was then processed for cGMP- and/or cAMP-PDE activity using a radiotracer assay based on ^32^, with some adjustments ^17, 18^. Briefly, samples were incubated with 35000-45000 disintegrations/minute of [^3^H]-cAMP or [^3^H]-cGMP for 10 minutes. The reaction was then quenched with 0.1M HCl and neutralized using 0.1M Tris. Snake venom was then added to the sample and incubated for 10 minutes at 37 °C. Samples were then run down DEAE A-25 Sephadex columns previously equilibrated in high salt buffer (20 mM Tris-HCl, 0.1% sodium azide, and 0.5M NaCl) and low salt buffer (20 mM Tris-HCl and 0.1% sodium azide). After washing the columns four times with 0.5 mL of low salt buffer, the eluate was mixed with 4 mL of scintillation cocktail, and then CPMs were read on a Beckman-Coulter liquid scintillation counter. 2 reactions not containing any sample lysate were also taken through the assay to assess background, which was subtracted from the sample CPMs. Data was expressed as CPMs/ug protein.

### Western Blotting

As previously reported^1^, cells were sonicated in ice cold lysis buffer (20 mM Tris-HCl, pH 7.5; 2 mM MgCl_2_; Thermo Pierce Scientific phosphatase tablet #A32959 and protease inhibitor 3 #P0044) for data in Figure 2C or in PDE assay buffer as described above. Protein concentrations were determined by DC Protein Assay kit (Bio-Rad; Hercules, CA, USA) according to manufacturer protocol, and were subsequently equalized across samples. Samples were stored at -80°C until further processing. For western blotting, 10μg of total protein was loaded onto 4-12% NuPAGE Bis-Tris gels (Invitrogen, Waltham MA) and electrophoresed for one hour at 180 volts. GFP-transfected cell samples were included on all PDE11A4 blots as a negative control. Protein was transferred onto a 0.45μm nitrocellulose membrane (Amersham, #10600008) for two hours at 100 mA. Membranes were then washed twice in tris-buffered saline with 0.1% tween20 (TBS-T) before staining with Ponceau S to determine total protein loading. Note, Ponceau S was chosen over a housekeeping gene as a loading control based on the best-practice statement of the Journal of Biological Chemistry ^33^. Images of the stained membranes were collected to later quantify the optical density of the total protein stain (i.e., spanning ∼200kDa to 10kDa), and then the membranes were rinsed in TBS-T to remove the stain. Blots to be probed with our custom PDE11A4 antibody (chicken polyclonal; Aves, #1-8113; 1:10,000) were blocked in 5% milk while those to be probed with GFP (rabbit polyclonal; Santa Cruz, #sc8334; 1:2000) were blocked in Superblock Blocking Buffer (ThermoFisher, Cat#37515), each with 0.1% Tween 20. Primary antibodies were incubated overnight at 4°C. The next day, membranes were washed 4 x 10 minutes with TBS-T and then incubated for 1 hour at room temperature with a secondary antibody (anti-chicken: Jackson Immunoresearch, 103-035-155 at 1:40 000; anti-rabbit: Jackson Immunoresearch, 111-035-144 at 1:10 000). Subsequently, membranes were washed 3 x 15 minutes in TBS-T. Finally, the membranes were immersed in SuperSignal West Pico Chemiluminescent Substrate as per manufacturer’s directions (ThermoScientific, Waltham MA), wrapped in clear a plastic sheet protector, and exposed to film. Multiple film exposures were taken to ensure signals were within the linear range, and Ponceau S stain and western blot optical densities were quantified using Image J. To account for membrane-membrane variances in film exposure, antibody saturation, chemiluminescence reaction, etc. between blots, Western blot data were normalized to a control condition (e.g. EmGFP-mPDE11A4 + vehicle) on each blot, as previously described (e.g., ^16, 34, 35^).

### Quantification of mPDE11A4 clustering into punctate spheroid shapes

As previously described ^15, 36^, all images pertaining to an experiment were quantified by an experimenter blind to treatment using the same computer within the same position in the room, the same lighting conditions, and the same percent zoom. Images were loaded onto a gridded template to facilitate keeping track of count locations within the image, and an experimenter scored each image box by box, with cells along the top and left edges of the entire image not included to follow stereological best practices. Images were quantified in a counterbalanced manner such that 1 picture from each condition was evaluated before moving onto a 2^nd^ image from that condition. The experimenter classified cells as exhibiting either only diffuse labeling or as having punctate structures (with/without diffuse labeling). Data are expressed as the % of the total number of labeled cells that exhibited punctate accumulations of the enzyme.

### In vivo study

As previously described^18^, young C57BL6 mice were imported from the National Institute of Aging (NIA) colony and were then bred onsite at the University of Maryland School of Medicine (UMSOM). *Pde11a* KO mice were also bred onsite at UMSOM. All mice were group-housed 4-5/cage, with young mice tested at ∼3 months old and old mice tested at ∼18 months old. Equal numbers of female and male mice were used.

As described by others^14^, we used peanut butter as a vehicle for oral delivery of SMQ-03-20. Briefly, Jif brand creamy peanut butter was melted to a liquid state in a sterile beaker by stirring it on a warming plate. The liquid peanut butter was either used plain or had a body weight-appropriate amount of SMQ-03-20 added to yield a 30 mg/kg dose to mice. The plain peanut butter was then slowly poured into molds using a 3 mL syringe without a needle to avoid bubbles or voids. The rectangular cavities of the mold hold 100 μL of liquid and a sterile razor blade was used to scrape off any excess peanut butter that escaped the cavity. The SMQ-03-20-laden peanut butter was poured similarly into separate SMQ-03-20-designated molds. Molds were then placed onto dry ice for ∼20 minutes to freeze and then were stored in an ultralow freezer until time of use. On the day of testing, pellets were removed from the freezer and immediately placed on dry ice where they remained until the precise time at which a mouse was dosed. Mice were food restricted for 1 night and the next morning (Day 1), all mice were transported to a quiet (∼50-52 dB), brightly lit room (∼700-800 lux) designated for *in vivo* studies. Mice were then singly housed in clean plastic cages with no bedding, allowed to habituate for 1 hour, and then provided a plain peanut butter pellet for habituation. Typically, mice took ∼10-60 minutes to eat the first pellet. At this point, ad lib food was returned to the home cage. On day 2, mice were again habituated to a plain peanut butter pellet, taking ∼7-8 minutes to consume the pellet. On day 3, half of the mice were provided a plain peanut butter pellet (vehicle) and the other half were given a peanut butter pellet containing a 30 mg/kg dose of SMQ-03-20. Generally, mice ate their pellets in ∼2-3 minutes.

2 hours after consuming their pellet, mice were moved to a second room and were killed by cervical dislocation. As previously described^1^, tissue was harvested fresh on wet ice, flash frozen in 2-methylbutane sitting on dry ice, placed directly on the dry ice to allow for evaporation of the 2-methylbutane. ½ brains were then stored long-term at -80°C until being processed via immunofluorescence.

### Immunofluorescence (IF)

Fresh-harvested, flash-frozen brains were embedded in matrix, cryosectioned in the sagittal plane at 20 μM, and thaw-mounted onto a +/+ slide. As previously described ^1, 30^, frozen tissue was fixed using 4% paraformaldehyde in 1X PBS for 15 minutes. After fixation, the tissue was washed 3 x 10 minutes in 1X PBS with bovine serum albumin and triton X-100 (PBT). Overnight incubation was conducted at 4°C using an antibody cocktail in PBT that included 2 antibodies that detect PDE11A4-pS117/pS124, a posttranslational modification found almost exclusively in clustered PDE11A4^1^ (Fabgennix PPD11A-140AP at 1:1000 and Fabgennix PPD11A-150AP at 1:500). The following day, sections were washed 4x in PBT and incubated for 90 min in secondary antibody (Alexafluor 488 AffiniPure Donkey Anti-Rabbit, 1:1000, Jackson Immunoresearch # 711-545-152). The secondary antibody was then washed off with PBT in 3X10 min washes. After rinsing the slides 10X in 1X PBS, slides were then treated with TrueBlack Lipofuscin Autofluorescence Quencher (Biotium, catalog # 23007). After, excess TrueBlack was rinsed off with agitation 3X in 1X PBS. Once dry, slides were mounted with DAPI fluoromount (Southern Biotech, #0100-20). Slides were kept covered and refrigerated until imaged using a Leica Model DM6 CFS microscope equipped with the Leica TCS SP8 laser system, Leica STP8000 control panel, Lumencor Sola Light Engine epifluorescent lamp, and Leica CTR6 power source. Representative images were captured with the Leica Application Suite X software using the HC PL APO 20x/0.74 IMM CORR CS2 objective with max projection images from Z-stacks saved as TIFF files.

### Counting ghost axons

Endogenous mPDE11A4 ghost axons in brain were quantified by an experimenter blind to treatment. PDE11A4 “ghost axons” are filamentous accumulations of PDE11A4. Ghost axons were quantified using the consistent application of classification standards with consideration to the continuity of the strand, quality of IF brightness, and proximity with other ghost axons. Non-specific or dull structures were omitted, while non-continuous strands of dots required further consideration in determining whether one or multiple ghost axons were present. Data accuracy was maintained by conducting each analysis under consistent room lighting, monitor settings, and time of day across all samples. Images were quantified using the “Analyze” function in the “ImageJ” application. The magnification was set to 75% and remained constant in all images. Data was gathered using the “Multi-Point” tool which incrementally recorded the number of ghost axons upon each click. The total number of ghost axons was then entered into an .xls spreadsheet for subsequent analysis.

### Data analyses

All between-group analyses were performed using Sigmaplot v11.2. Age and/or treatment effects were analyzed by One-way or Two-way ANOVA (F) when normality as per the Shapiro-Wilks test and/or equal variance as per the Levene’s test passed. When these assumptions were not met, then an ANOVA on ranks (H) or Mann Whitney Rank Sum Test (T) was used. Following a significant main effect or interaction, *post hoc* tests were conducted using Student Newman-Keuls method and significance was defined as P<0.05. Please note that Sigmaplot provides exact P-values for *post hoc* tests following parametric ANOVAs, but only yes or no to “P<0.05” for *post hoc* tests following a significant non-parametric ANOVA. Data are graphed as mean ± standard error of the mean (SEM).

## RESULTS

### PDE11A4 exhibits sequence features associated with liquid:liquid phase separation (LLPS)

Only a subset of protein sequences are capable of undergoing LLPS under physiological conditions, and they generally involve multivalent sequences that enable protein interactions with a scaffolding protein (i.e., heterotypic condensation) or within itself (i.e., homotypic condensation; c.f., ^22^). These “pro-LLPS” protein sequences include IDRs such as the one in the N-terminal of PDE11A4; however, not all IDRs promote LLPS^223^. IDRs that promote LLPS typically contain additional features including phosphorylation and/or other post-translational modifications, non-covalent pi-pi interactions, prion-like domains, low-complexity regions, and a certain fraction and patterning of charged residues (c.f., ^22, 27, 28^). As such, we utilized several recommended computational screening tools^22, 27, 28^ to determine if 1) PDE11A4 is likely to have additional IDRs outside of the N-terminal and 2) if any IDRs exhibit LLPS-promoting features other than the previously described pS117/pS124 motif that falls within the N-terminal IDR^1^. D2P2, a program that synthesizes analyses provided by 9 different disorder prediction algorithms^2^, identifies several IDRs within PDE11A4 that are conserved across human and mouse (Figure 1). That said, only the annotated N-terminal IDR is enriched in ANCHOR domains^37, 38^. That is, regions that do not form favorable interactions that sustain stable structures on their own but are capable of sustaining a stable structure when interacting with a globular structure (i.e., a disorder-to-order transition). Analyses for non-covalent pi-pi interactions using PScore^6^ and prion-like domains using PLAAC^7^ identified a conserved enrichment for both motifs within PDE11A4, but again only within the N-terminal IDR (Figure 1). Analysis for low-complexity regions by SEG^8^, as reported by PhaSePred^5^, reveals 2 low-complexity regions in the N-terminal IDR of both human and mPDE11A4 and 1 in the C-terminal IDR of hPDE11A4 only.

**Figure 1.**
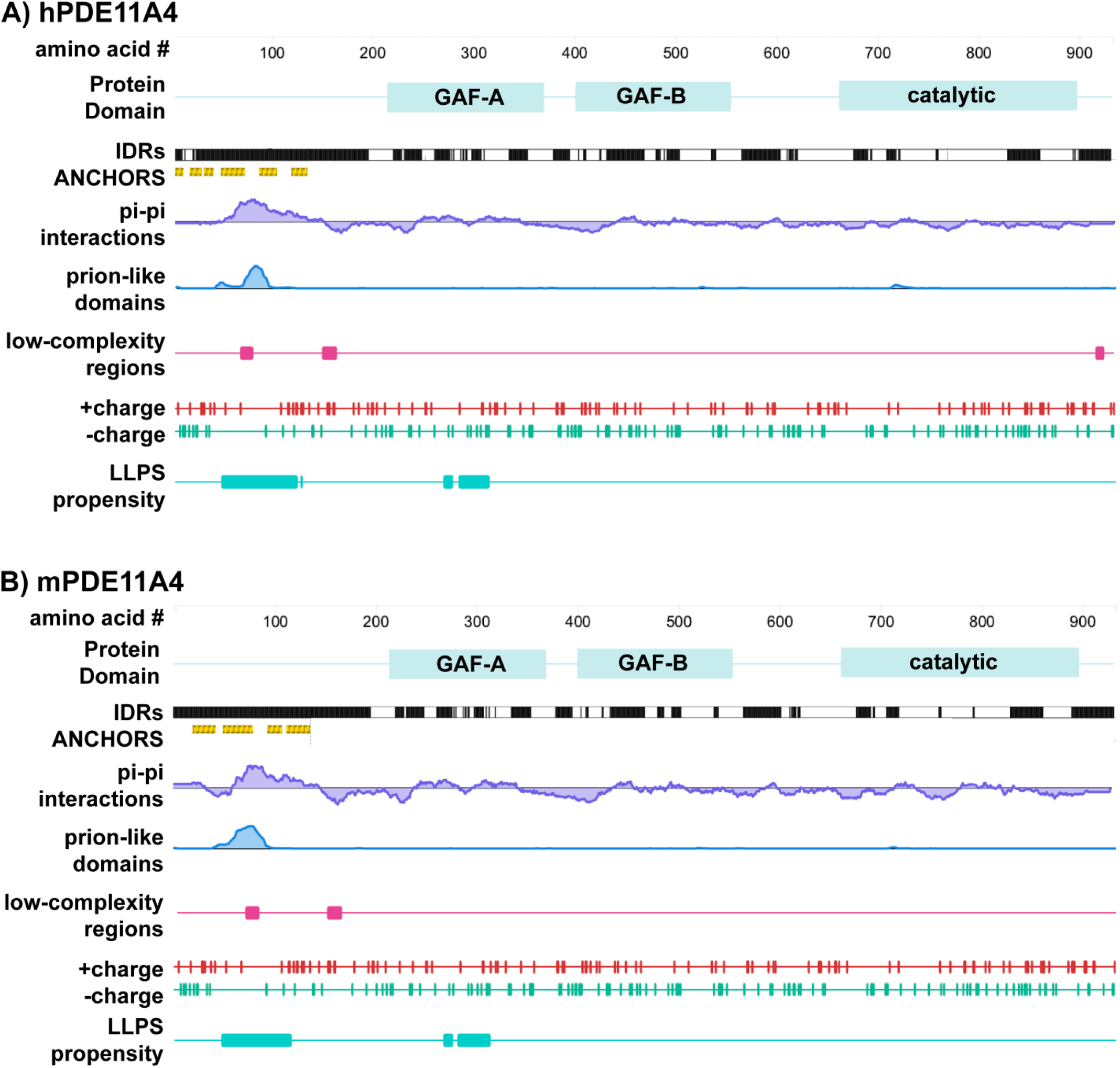
Multiple computational analyses identify LLPS-promoting sequence features within PDE11A4. Graphical outputs from D2P2^2^ (IDRs and ANCHORS) and PhaSePred^5^ (all other outputs) for A) human PDE11A4 (hPDE11A4) and B) mouse PDE11A4 (mPDE11A4). PhaSePred reported analyses from other primary computational tools as follows: pi-pi from Pscore^6^, prion-like domain from PLAAC^7^, low-complexity regions from SEG^8^, and LLPS propensity from catGRANULE^9^. GAF--mammalian c<u>G</u>MP-regulated PDEs, Anabaena <u>a</u>denylyl cyclases, and the Escherichia coli transcription factor <u>F</u>hlA; IDRs—intrinsically disordered regions.

Calculation of the distribution and patterning of charged and hydrophobic residues by CIDER^39^ places both human and mouse PDE11A4 in the “R1” category since they are weak polyampholytes (0.1147 and 0.1136, respectively) and weak polyelectrolytes (0.1276 and 0.1297, respectively). Thus, they are predicted to form globules (see Figure S1 for the Das Pappu Diagram). Interestingly, CIDER analyses suggest the formation of these PDE11A4 globules may be context dependent given the proximity of its score to the R2 category and the fact that their fraction of charged residues (FCR) is 0.2422 and 0.2243, respectively (i.e., just below the 0.25 cutoff to be considered R2). Consistent with the fact that the fraction of positively charged residues (*f*+) is approximately equal to the fraction of negatively charged residues (*f-*), the “net charge per residue” (NCPR) score is close to zero at -0.0129 and -0.016, respectively, suggesting PDE11A4 may behave as disordered globules that are regulated by attractive interactions^24, 39^. Finally, CIDER calculates the charge patterning κ scores for human and mPDE11A4 as 0.1727 and 0.1736, respectively, both of which fit comfortably within the range of 0.1-0.3 that is highly predictive of proteins undergoing LLPS^24, 39^.

In addition to the above algorithms that identify individual LLPS-promoting features, there are additional tools that synthesize the output from several of these tools to calculate an overall likelihood of undergoing LLPS. One such tool is catGRANULE^9^, an algorithm that calculates the LLPS propensity of each residue in a given sequence by combining structural disorder, sequence length, nucleic acid binding propensities, and the content of glycine, arginine, and pheylalanine^27^. The overall catGRANULE scores for human and mPDE11A4 as reported by PhaSePred^5^ are 0.481 and 0.420, respectively. Interestingly, catGRANULE predicts high LLPS propensity for regions of the protein that fall within the N-terminal IDR and the regulatory GAF-A domain (Figure 1). PhaSePred^5^ does not simply report catGRANULE, Pscore, PLAAC, IDR, hydropathy, low-complexity, and charged residues scores all in one place, it also ranks those individual scores amongst all proteins in that species and generates a synthesized ranking for likelihood of undergoing LLPS (0 lowest-1 highest). Strikingly, PhaSePred^5^ ranks the Pscore and PLAAC scores for hPDE11A4 at 0.924 and 0.913 out of 1.0, respectively. Further, PhaSePred ranks PDE11A4 as likely to form condensates via both self-assembly (hPDE11A4: score 0.34, rank 0.74; mPDE11A4: score 0.215, rank .7) and via interactions with other protein and/or nucleic acid binding partners (hPDE11A4: score 0.46, rank 0.69/1; mPDE11A4: score .53, rank 0.73/1). Phase Separation Predictor ^40^ agrees that both hPDE11A4 and mPDE11A4 will phase separate, each with a score of 0.8861 out of 1.0.

### PDE11A4 exhibits physical properties associated with LLPS

Given that a variety of bioinformatic tools predict that both mouse and hPDE11A4 undergo LLPS, we next determined if PDE11A4 demonstrated physical properties of LLPS. At a minimum, proteins that undergo LLPS should form membraneless spherical droplets that fuse over time in a concentration dependent manner and are reversible ^23, 25^. Since manipulations of LLPS would be functionally explored *in vivo* in the mouse hippocampus, we focused our *in vitro* studies using mPDE11A4. To determine if PDE11A4 meets these 4 criteria, we transfected HT22 or COS1 cells with mPDE11A4 that was N-terminally fused with EmGFP to facilitate microscopic detection. It is highly unlikely that the EmGFP tag is driving the clustering of mPDE11A4 since 1) EmGFP alone does not form spherical clusters even when expressed at very high levels (Figure 2C), 2) immunocytochemistry shows untagged hPDE11A4 similarly forms spherical clusters *in vitro*^1, 36^, 3) EmGFP-mPDE11A4 retains its catalytic activity and proper subcellular compartmentalization upon biochemical fractionation^1^, and 4) we introduced an A206Y point mutation into our EmGFP tag to prevent dimerization ^29^. We first used EmGFP-mPDE11A4 to assess if spherical mPDE11A4 droplets are free of membranes. To do so, we applied BODIPY-568 to label membranes of mouse hippocampal HT22 cells transfected with mPDE11A4. Consistent with our previously described inability to colocalize mPDE11A4 spherical droplets with standard organelle markers^1^, we could find no evidence of colocalization of the mPDE11A4 spherical droplets with a membrane signal across 4 coverslips (Figure 2A).

**Figure 2.**
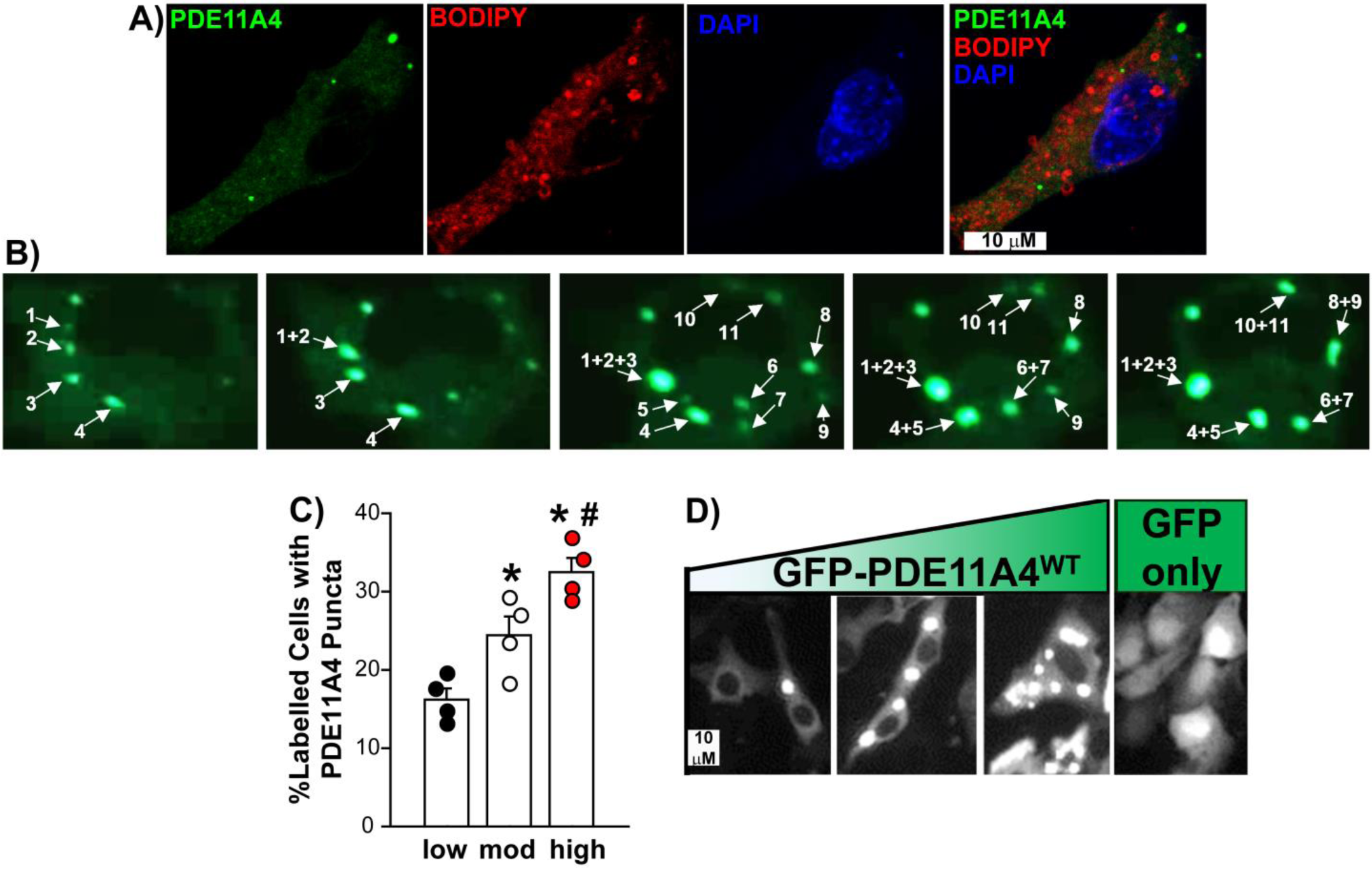
mPDE11A4 forms membraneless spherical droplets that fuse over time in a concentration dependent manner. A) Membrane staining using BODIPY-568 in HT22 cells transfected with EmGFP-mPDE11A4 shows no co-localization of the signals (representative image of cells across 4 coverslips). B) Time-lapsed imaging of COS1 cells expressing EmGFP-mPDE11A4 over 120 minutes clearly shows droplets fusing over time (representative of 10 cells over 3 experiments). C) Quantification and D) representative images of HT22 cells expressing low, <u>mod</u>erate, or high levels of mPDE11A4 protein show mPDE11A4 droplets form in a concentration-dependent manner (n=4 biological replicates/group). Also GFP alone shows no punctate clustering despite high expression throughout the cell, particularly the nucleus. *vs low, P=0.0146-0.0007, #vs high, P=0.0158. mod—moderate, GFP—emerald green fluorescent protein.

Second, we determined if the spherical mPDE11A4 droplets fuse over time. Since our previous studies showed that both HT22 and COS1 cells similarly form mouse and human PDE11A4 spherical droplets^1^, here we used COS1 cells as their larger size makes it easier to visualize and track the mPDE11A4 droplets over time. Indeed, time-lapsed imaging of 10 cells over 3 experiments demonstrates that mPDE11A4 spherical droplets emerge over time following a transient transfection and eventually fuse with each other (Figure 2B). Smaller mPDE11A4 droplets were also noted to separate into even smaller droplets on rare occasions. In contrast, larger mPDE11A4 spherical droplets appeared to become less dynamic over time and could adopt an “irregular shape” that may reflect a liquid:gel phase transition^22^.

Finally, we established if mPDE11A4 droplets form in a concentration-dependent manner. To do so, we compared mPDE11A4 droplet formation in HT22 cells transfected with a low (0.0375 μg cDNA/mL of media), moderate (0.1125 μg cDNA/mL of media), or high concentration of plasmid (0.375 μg cDNA/mL of media).

First, we verified that such titration of cDNA yielded low, moderate, versus high levels of mPDE11A4 protein expression levels by Western blot (Table 1; equal variance failed; ANOVA on Ranks: H(2)=9.85, P=0.0002; *Post hoc*: each vs the other, P<0.05). Next, we determined if these varying levels of mPDE11A4 protein would correspond to increasing levels of mPDE11A4 clustering. Indeed, the presence of mPDE11A4 LLPS-like droplets significantly increased as protein levels increased (F(2,9)=17.96, P=0.0007; Post hoc: low vs. moderate P=0.0146, moderate vs. high P=0.0158). Together, these data show that PDE11A4 meets 3 of the 4 minimal requirements for LLPS in that it forms membraneless spherical droplets that fuse over time in a concentration dependent manner (see more below for 4^th^ requirement—reversibility).

**Table 1.** mPDE11A4 protein expression (mean ±SEM) in HT22 cells transfected with differing concentrations of EmGFP-mPDE11A4 plasmid (μg plasmid DNA/mL of media).

| **Table 1.** mPDE11A4 protein expression (mean ±SEM) in HT22 cells transfected with differing concentrations of EmGFP-mPDE11A4 plasmid (µg plasmid DNA/mL of media). | |
| --- | --- |
| **Amount of plasmid** | **mPDE11A4/Ponceau**<br>**r.o.d.** |
| Low (0.0375 µg/mL) | 0.06 ±0.04 |
| Moderate (0.112 µg/mL) | 0.72 ±0.2 |
| High (0.375 µg/mL) | 3.82 ±0.85 |

### PDE11Ai’s inhibit mouse PDE11A4 catalytic activity in mouse HT22 hippocampal cells

Here we report findings with 5 PDE11i inhibitors (PDE11i’s) across 3 scaffolds. The first scaffold is represented by Tadalafil (Figure 3A), a PDE5A inhibitor approved for treatment of erectile dysfunction and benign prostatic hypertrophy that is well known to potently inhibit PDE11A4^41^. The second scaffold is represented by BC11-38 (Figure 3B), a selective PDE11Ai tool compound previously identified by the Hoffman lab ^42^. The third scaffold is represented by three “SMQ” compounds previously reported to exhibit potent, selective inhibition of PDE11A4 *in vitro* in cells (Figure 3C-D) ^17, 18^. The biochemical IC_50_ values against human PDE11A4 for these compounds are as follows: tadalafil 25 nM^17^, BC11-38 280 nM^42^, SMQ-2-057 12 nM^17^, SMQ-3-030 34 nM^18^, SMQ-3-020 15 nM^18^.

**Figure 3.**
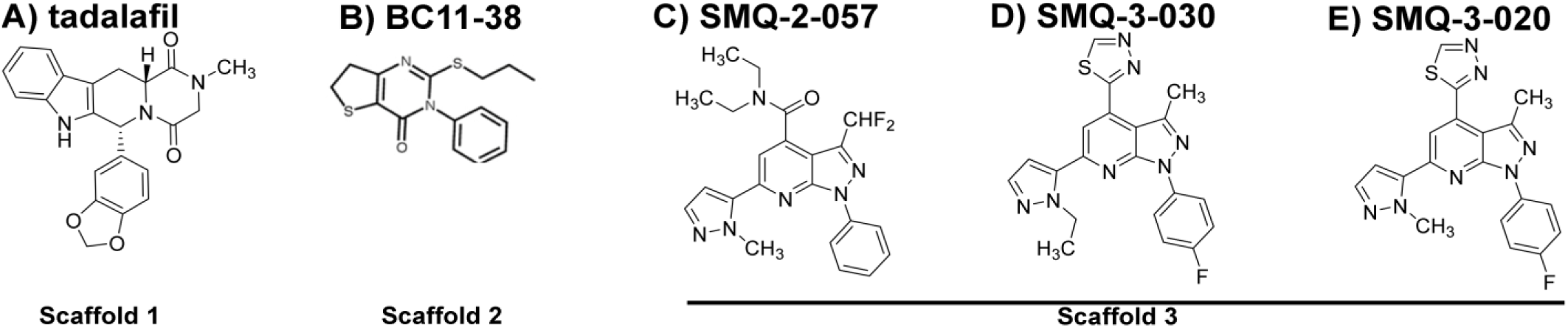
Chemical structures for A) tadalafil, B) BC11-38, C) SMQ-02-057, D) SMQ-03-30, E) SMQ-03-20.

Before testing effects of these inhibitors on mouse PDE11A4 subcellular compartmentalization, we first verified that these PDE11Ai’s identified in screens against human PDE11A4 do inhibit mouse PDE11A4 (mPDE11A4) catalytic activity in mouse HT22 hippocampal cells (Table 1). Please note that the PDE activity reported in Table 1 for tadalafil, SMQ-02-57, and SMQ-03-20 were previously reported in graphical format^17, 18^. Data for BC11-38 and SMQ-03-30 are presented here for the first time. Notably, both cAMP- and cGMP-PDE activity is very low in GFP-transfected HT22 cells treated with vehicle (i.e., GFP + 0 μM) but is greatly increased in vehicle-treated cells expressing mPDE11A4 (i.e., WT + 0 μM). This mPDE11A4-mediated *cAMP-PDE* activity was significantly inhibited by <u>10-100 μM tadalafil</u> (as originally reported for “1” in^17^; fails normality; ANOVA on Ranks: H(5)=19.01, P=0.0019; Post hoc: WT + 0 μM vs WT + 10 μM-100 μM, P<0.05 for each), <u>100 μM BC11-38</u> (F(5,18)=76.79, P<0.0001; Post hoc: WT + 0 μM vs. WT + 100 μM P=0.0004), <u>1-100 μM SMQ-02-57</u> (as originally reported for “23b” in^17^; normality fails; ANOVA on Ranks: H(5)=21.48, P=0.0007; Post hoc: WT + 0 μM vs. WT + 1-100 μM, P<0.05 for each), <u>1-100 μM SMQ-03-30</u> (reported as “4h” in^18^; F(5,18)=85.79, P<0.0001; Post hoc vs. WT + 0 μM: WT + 1 μM P=0.0019, WT + 10 μM P=0.0001, WT + 100 μM P=0.0002), and <u>1-100 μM SMQ-03-20</u> (as originally reported for “4g” in^18^; fails equal variance; ANOVA on Ranks: H(5)=21.77, P=0.0006; Post hoc: WT + 0 μM vs WT + 1-100 μM, P<0.05 for each). Similarly, mPDE11A4 cGMP-PDE activity was significantly inhibited by <u>100 μM tadalafil</u> (fails normality; ANOVA on Ranks: H(5)=19.13, P=0.0018; *Post hoc:* WT + 0 μM vs WT + 100 μM, P<0.05), <u>100 μM BC11-38</u> (Figure 1L; F(5,18)=59.93, P<0.0001; Post hoc: WT + 0 μM: vs. WT + 100 μM P=0.0019), <u>0.1-100 μM SMQ-02-57</u> (Figure 1M; normality fails; ANOVA on Ranks: H(5)=21.45, P=0.0007; Post hoc: WT + 0 μM vs. WT + 0.1-100 μM, P<0.05 for each), <u>10-100 μM SMQ-03-30</u> (Figure 1N; equal variance failed; ANOVA on Ranks: H(5)=20.39, P=0.001; Post hoc: WT + 0 μM vs WT + 10-100 μM, P<0.05 for each), and <u>10-100 μM SMQ-03-20</u> (Figure 1O; F(5,18)=451.34, P<0.0001; Post hoc vs WT + 0 μM: WT + 1 μM P=0.0792, WT + 10 μM P=0.0002, WT + 100 μM P=0.0001).

**Table 1.**
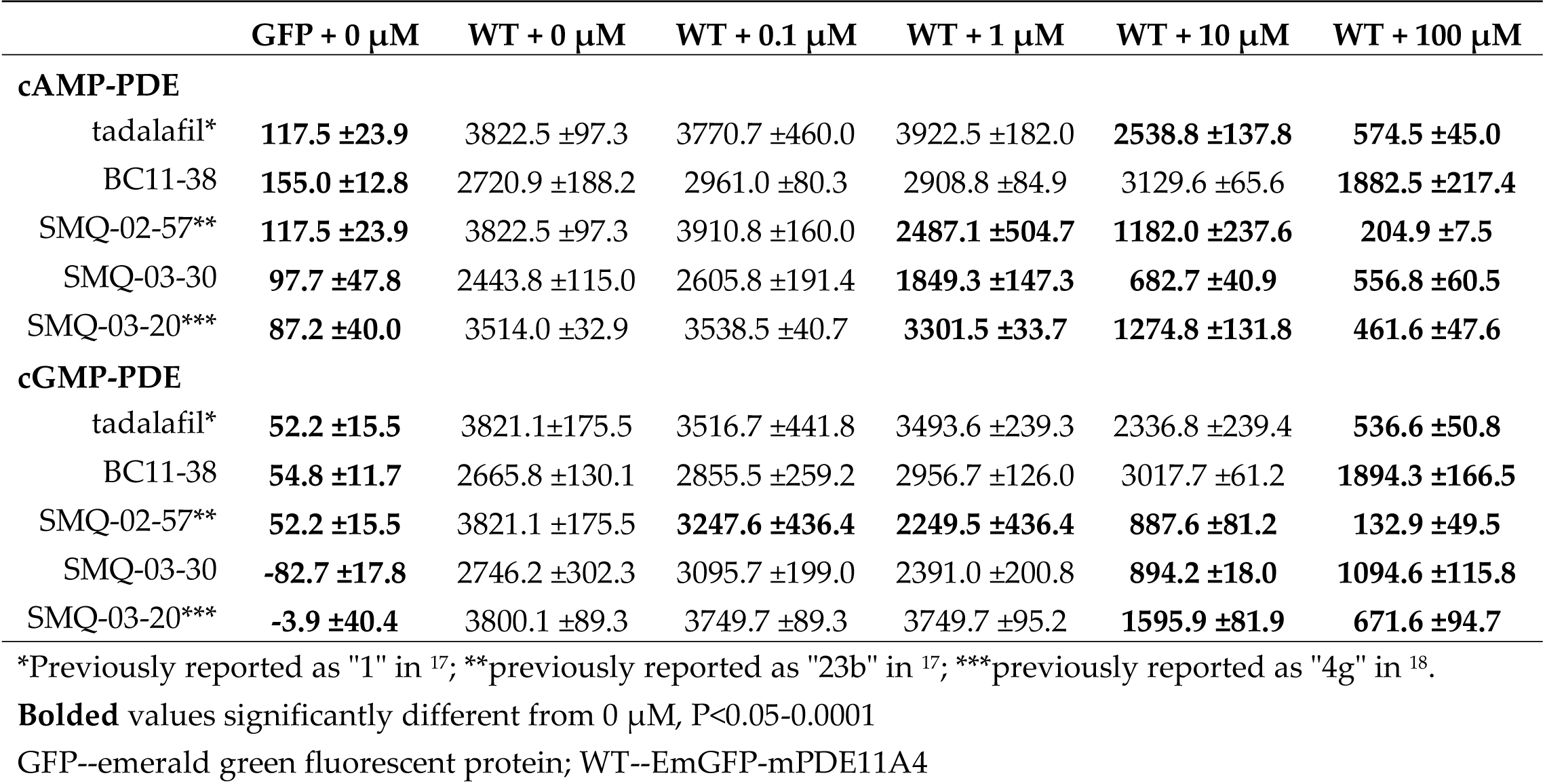
PDE11A inhibitors identified in human screen also inhibit mouse PDE11A4 cAMP- and cGMP-catalytic activity (expressed as the mean CPMs/μg protein ±SEM) in mouse HT22 hippocampal cells.

| | GFP + 0 μM | WT + 0 μM | WT + 0.1 μM | WT + 1 μM | WT + 10 μM | WT + 100 μM |
| --- | --- | --- | --- | --- | --- | --- |
| **cAMP-PDE** |  |  |  |  |  |  |
| tadalafil* | **117.5 ±23.9** | 3822.5 ±97.3 | 3770.7 ±460.0 | 3922.5 ±182.0 | **2538.8 ±137.8** | **574.5 ±45.0** |
| BC11-38 | **155.0 ±12.8** | 2720.9 ±188.2 | 2961.0 ±80.3 | 2908.8 ±84.9 | 3129.6 ±65.6 | **1882.5 ±217.4** |
| SMQ-02-57** | **117.5 ±23.9** | 3822.5 ±97.3 | 3910.8 ±160.0 | **2487.1 ±504.7** | **1182.0 ±237.6** | **204.9 ±7.5** |
| SMQ-03-30 | **97.7 ±47.8** | 2443.8 ±115.0 | 2605.8 ±191.4 | **1849.3 ±147.3** | **682.7 ±40.9** | **556.8 ±60.5** |
| SMQ-03-20*** | **87.2 ±40.0** | 3514.0 ±32.9 | 3538.5 ±40.7 | **3301.5 ±33.7** | **1274.8 ±131.8** | **461.6 ±47.6** |
| **cGMP-PDE** |  |  |  |  |  |  |
| tadalafil* | **52.2 ±15.5** | 3821.1 ±175.5 | 3516.7 ±441.8 | 3493.6 ±239.3 | 2336.8 ±239.4 | **536.6 ±50.8** |
| BC11-38 | **54.8 ±11.7** | 2665.8 ±130.1 | 2855.5 ±259.2 | 2956.7 ±126.0 | 3017.7 ±61.2 | **1894.3 ±166.5** |
| SMQ-02-57** | **52.2 ±15.5** | 3821.1 ±175.5 | **3247.6 ±436.4** | **2249.5 ±436.4** | **887.6 ±81.2** | **132.9 ±49.5** |
| SMQ-03-30 | **-82.7 ±17.8** | 2746.2 ±302.3 | 3095.7 ±199.0 | 2391.0 ±200.8 | **894.2 ±18.0** | **1094.6 ±115.8** |
| SMQ-03-20*** | **-3.9 ±40.4** | 3800.1 ±89.3 | 3749.7 ±89.3 | 3749.7 ±95.2 | **1595.9 ±81.9** | **671.6 ±94.7** |

\*Previously reported as "1" in <sup>17</sup>; \*\*previously reported as "23b" in <sup>17</sup>; \*\*\*previously reported as "4g" in <sup>18</sup>.
**Bolded** values significantly different from 0 μM, $P<0.05$ -0.0001
GFP--emerald green fluorescent protein; WT--EmGFP-mPDE11A4

Importantly, these reductions in catalytic activity occurred in the absence of any significant changes in mPDE11A4 protein expression (Table 2). Together, these results confirm that these compounds inhibit mouse PDE11A4 (mPDE11A4) as well as human PDE11A4.

**Table 2.**
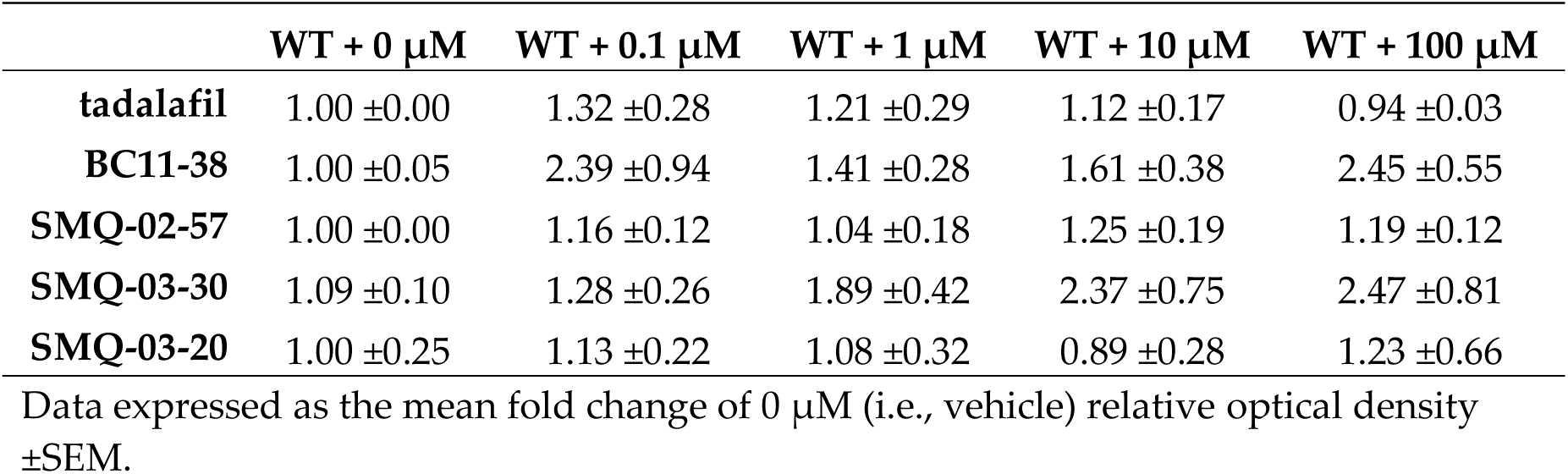
PDE11A inhibitors do not change protein expression of PDE11A4 in HT22 hippocampal cells.

### PDE11Ai’s reverse aging-like clustering of mPDE11A4 protein in mouse HT22 hippocampal cells

As described above, age-related increases in mPDE11A4 protein expression trigger ectopic clustering of the enzyme in “ghost axons” in brain, and this phenomenon can be modeled in HT22 cells^1^ (Figure 2). Thus, we next determined if PDE11Ai’s were capable of reversing this age-related clustering of mPDE11A4 in HT22 cells (Figure 4). Strikingly, all 5 PDE11A4i’s dramatically reverse aging-like clustering of mPDE11A4 in HT22 cells in less than 1 hour (Figure 4). Relative to vehicle-treatment (i.e., 0 μM), the percentage of cells exhibiting punctate clustering of mPDE11A4 was reduced by <u>10-100 μM tadalafil</u> (Figure 4B; F(4,15)=32.44, P<0.0001; Post hoc: WT + 0 μM vs WT + 10-100 μM, P=0.0002 for each), <u>100 μM BC11-38</u> (Figure 4C; F(), <u>0.1-100 μM SMQ-02-57</u> (Figure 4D; equal variance fails; ANOVA on Ranks: H(4)=15.61, P=0.0036; Post hoc: WT + 0 μM vs. WT + 0.1-100 μM, P<0.05 for each), <u>1-100 μM SMQ-03-30</u> (Figure 4E; equal variance fails; ANOVA on Ranks: H(4)=16.74, P=0.0022; Post hoc:s. WT + 0 μM vs WT + 1-100 μM, P<0.05 for each), and <u>1-100 μM SMQ-03-20</u> (Figure 4F; fails equal variance; ANOVA on Ranks: H(4)=17.43, P=0.0016; Post hoc: WT + 0 μM vs WT + 1-100 μM, P<0.05 for each). Importantly, this dispersal of mPDE11A4 clusters by PDE11Ai’s appears to be specific in that neither the PDE4 inhibitor rolipram nor the PDE10 inhibitor rolipram elicited such an effect (Figure 4G-H). Thus, across scaffolds, PDE11Ai’s reverse LLPS-like clustering of PDE11A4 protein in hippocampal HT22 cells. As such, PDE11A4 meets the 4^th^ out of 4 minimal requirements for LLPS—that the condensates are reversible.

**Figure 4.**
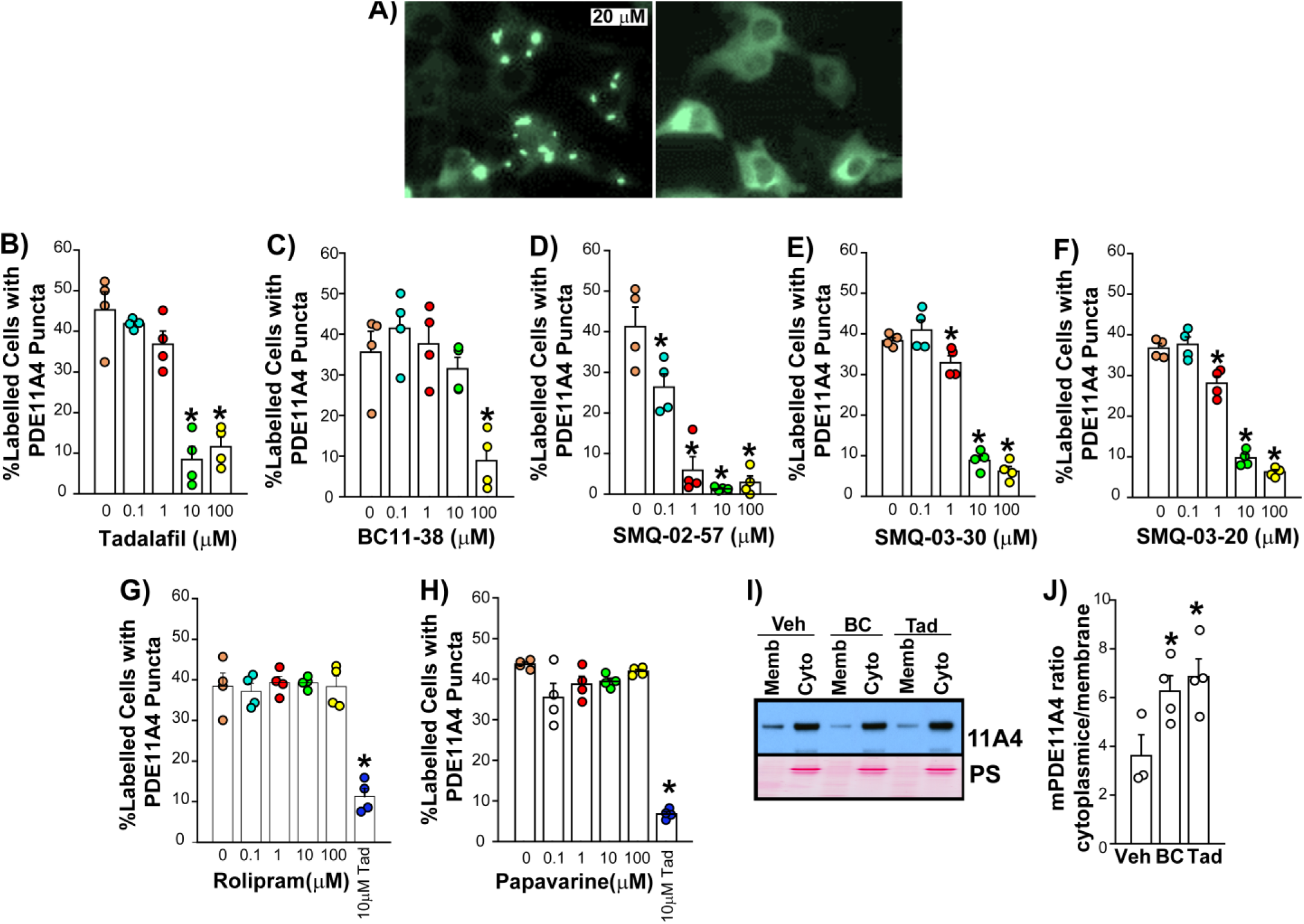
Five PDE11A4 inhibitors from 3 different scaffolds all reverse age-related clustering of mPDE11A4 in mouse hippocampal HT22 cells. A) Exemplar image of aging-like clusters of EmGFP-mPDE11A4 in HT22 cells following vehicle treatment (left) and their subsequent dispersal following treatment with a PDE11A4 inhibitor (right; shown: 1 μM SMQ-02-57). All PDE11A4 inhibitors tested to date reverse aging-like clustering of mPDE11A4 in a concentration-dependent manner, including B) tadalafil, C) BC11-38, D) SMQ-02-057, E) SMQ-03-30, AND F) SMQ-03-20. This dispersing effect is selective for PDE11A inhibitors in that neither G) the PDE4 inhibitor rolipram nor H) the PDE10 inhibitor papaverine reversed aging-like clustering of mPDE11A4 (10 μM <u>tad</u>alafil included as positive control). I) Representative western blot (top: EmGFP-PDE11A4; bottom: Ponceau Stain (PS)) and I) quantification showing aging-like accumulation of mPDE11A4 in the membrane vs. cytosolic fraction that is reversed by BC11-38 (BC) and Tadalafil (Tad; i.e. PDE11Ai’s shift mPDE11A4 from the membrane to the cytosol). *vs 0 μM P<0.05-0.0001.

### PDE11Ai’s shift mPDE11A4 from the membrane to the cytosolic fraction

Since PDE11Ai’s so convincingly reversed aging-like clustering of mPDE11A4, we next determined if PDE11Ai’s would also affect the distribution of mPDE11A4 between the cytosolic versus membrane compartments. Following biochemical fractionation, PDE11A4 protein is expressed at much higher levels in the cytosolic (a.k.a. soluble) fraction relative to the membrane (a.k.a. particulate) fraction in both in the hippocampus and HT22 cells^1^. Interestingly, age-related increases in ventral hippocampal mPDE11A4 are localized to the membrane fraction^1^, thus decreasing the ratio of cytoplasmic:membrane mPDE11A4. We see here that a 1-hour treatment of HT22 cells with 100 μM of either tadalafil (n=4, 6.26 ±0.64 A.U.) or BC11-38 (n=4, 6.86 ±0.73 A.U.) had the opposite effect of aging by reducing membrane mPDE11A4 thereby increasing the mPDE11A4 cytoplasmic:membrane ratio relative to vehicle (n=3, 3.61 ±0.86 A.U.; F(2,8)=4.99, P=0.039; Post hoc vs. vehicle: tadalafil P=0.0391, BC11-38 P=0.0389; Figure 4I-J;).

### Aging-like clusters of mPDE11A4 can reform following PDE11Ai washout

Next, we determined if mPDE11A4 aging-like clusters would reform following washout of the PDE11Ai’s, and if this process would be impacted by the duration of treatment or concentration of compound used. To do so, mPDE11A4-transfected HT22 cells were treated with vehicle or a PDE11Ai (i.e., tadalafil, BC11-38, or SMQ-02-57) for either 1 or 24 hours and were then either fixed immediately or returned to media without compound for a 5 hour-washout period and then fixed (Figure 5A). Tadalafil reversed aging-like mPDE11A4 clustering to a similar extent with either a 1-hour (F(4,15)=32.44, P<0.0001) or 24-hour treatment (F(4,15)=28.10, P<0.0001). With just a 5-hour washout of the drug, mPDE11A4 re-clustered (a.k.a. demixed) in the cells treated with 10 μM tadalafil for either 1 hour (F(4,15)=22.75, P<0.0001) or 24 hours (fails equal variance; ANOVA on Ranks: H(4)=15.16, P=0.0044), but not in cells treated with 100 μM (Post hoc: 1-hour WT + 0 μM vs. WT + 100 μM, P=0.0002; 1-hour WT + 10 μM vs. WT + 100 μM, P=0.0002; 24-hour WT + 0 μM vs. WT + 10-100 μM, P<0.05; 24-hour WT + 10 μM vs. WT + 100 μM, P<0.05). BC11-38 equivalently reduced mPDE11A4 clustering with either a 1-hour (F(4,15)=5.29, P=0.0073) or 24-hour treatment of 100 μM (F(4,15)=11.72, P=0.0002). Following a 5-hour washout, no significant effect of either the 1-hour (equal variance failed; ANOVA on Ranks: H(4)=7.99, P=0.092) or 24-hour treatment with 100 uM BC11-38 remained (F(4,15)=2.75, P=0.0673). SMQ-02-57 also reversed aging-like mPDE11A4 clustering to a similar extent with either a 1-hour (equal variance fails; ANOVA on Ranks: H(4)=16.0, P=0.0023; Post hoc vs WT + 0 μM: WT + 1-100 μM P<0.05 for each) or 24-hour treatment (F(4,15)=76.29, P<0.0001; Post hoc vs WT + 0 μM: WT + 0.1 μM P=0.0268, WT + 1 μM P=0.0004, WT + 10 μM P=0.0002, WT + 100 μM P=0.0001). Similarly to tadalafil, a 5-hour washout period was sufficient for mPDE11A4 clusters to reform following a 1-hour (F(4,15)=61.43, P<0.0001) or 24-hour treatment (F(4,15)=111.29, P.0.0001) with low but not high concentrations of SMQ-02-57 (1-hour Post hoc vs. WT + 0 μM: WT + 0. 1 μM P=0.1755, WT + 1 μM P=0.2938, WT + 10 μM P=0.0002, WT + 100 μM P=0.0001; 24-hour Post hoc vs WT + 0 μM: WT + 0. 1 μM P=0.8258, WT + 1 μM P=0.463, WT + 10 μM P=0.0002, WT + 100 μM P=0.0002). A follow-up study showed that the percentage of labelled cells exhibiting mPDE11A clusters significantly increases with just a 1-hr washout of 1 μM SMQ-02-57 (Table 3; failed equal variance, ANOVA on Ranks: H(2)=9.84, P=0.0002; *Post hoc*: each group vs. the other, P<0.05).

**Figure 5.**
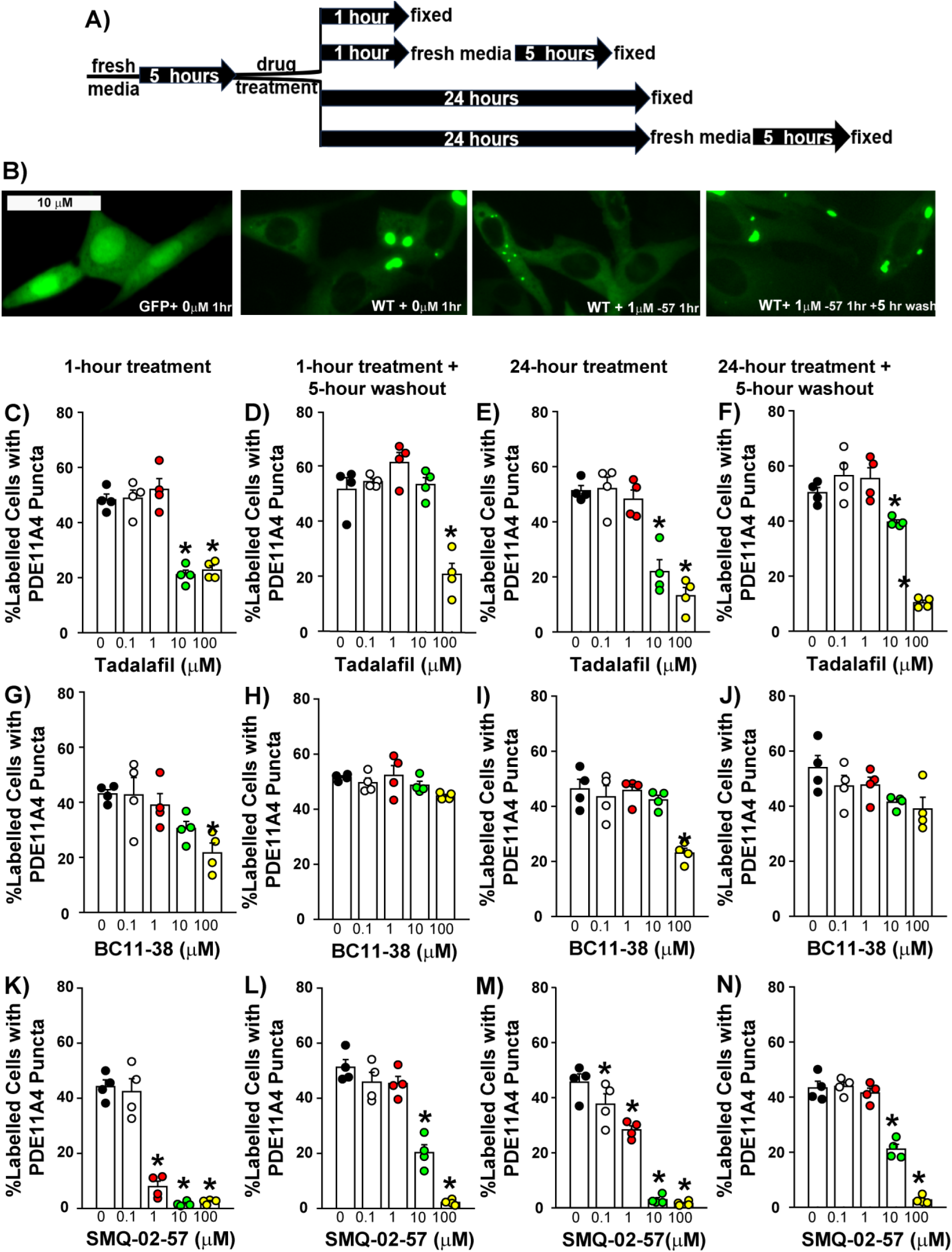
Dispersing effects of PDE11A4 inhibitors do not diminish with a 24-hour treatment and effects of lower concentrations are reversible upon washout. A) Graphical outline of experimental time course. B) Exemplar image of aging-like clusters of EmGFP-mPDE11A4 following vehicle treatment (left), their subsequent dispersal following treatment with a PDE11A inhibitor (middle; shown: 1 μM SMQ-02-57), and their reformation following washout of the compound for 1 or 5 hours (right). Extent of mPDE11A4 clustering measured following a 1-hour treatment, a 1-hour treatment followed by a 5-hour washout period, a 24-hour treatment and a 24-hour treatment followed by a 5-hour washout period for C-F) tadalafil, G-J) BC11-38, and K-N) SMQ-02-057. *vs 0 μM P<0.05-0.0001.

**Table 3.** mPDE11A4 clusters reform with only a 1-hr wash out following a 1-hr treatment with 1 μM SMQ-02-57 (mean ±SEM), but they appear to be smaller in size and in greater numbers relative to vehicle-treatment.

| **group** | **% Labelled cells**<br>**with puncta** | **#puncta/puncta-**<br>**containing cell** |
| --- | --- | --- |
| WT + 0 μM | 70.2 ±5.1 | 2.2 ±0.1 |
| WT + 1 μM SMQ-02-57 | 34.1 ±3.4 | 5.7 ±0.9 |
| WT + 1 μM SMQ-02-57 +<br>1-hr wash | 53.8 ±1.9 | 4.0 ±0.5 |

Qualitatively, it was noticed that the clusters in the 1-hour washout group appeared to be smaller in size and greater in number than we typically observe in the vehicle treatment or 5-hour washout groups. Therefore, we conducted a *post hoc* analysis of the average number of puncta/puncta-containing cell. Indeed, the number of puncta/puncta-containing cell was significantly greater in the 1 μM SMQ-02-57 treatment group and the 1-hr washout group relative to vehicle (failed equal variance, ANOVA on Ranks: H(2)=8.35, P=0.002; Post hoc vs. 0 μM: each P<0.05.

### Oral dosing of SMQ-03-20 to old NIA C57BL6 mice reverses age-related clustering of PDE11A4 in ghost axons

Finally, we attempted to confirm that oral administration of a brain-penetrant PDE11A4 inhibitor would be able to reverse age-related clustering of PDE11A4 *in vivo*. For this study we selected SMQ-03-20 previously shown to have improved brain penetration following oral dosing relative to tadalafil ^18, 43^. SMQ-03-20 is also sufficiently selective versus other PDE families, ∼34% bioavailability, and has a plasma and brain T1/2 of ∼1.5hr as well as a brain Tmax of 2 hours ^18^. Note that *in vivo* dosing of this compound is limited to 30 mg/kg due to solubility issues^18^. To capture effects on both catalytic activity and subcellular compartmentalization, we harvested tissue 2 hours after oral dosing of either peanut butter alone (i.e., vehicle) or peanut butter pellets laden with a 30 mg/kg dose of SMQ-03-20. Previously, we reported that oral dosing of 30 mg/kg significantly inhibited cAMP-PDE activity in the brain of young and old female and male NIA C57BL6 mice but not *Pde11a* KO mice, illustrating specificity of the compound for PDE11A4 in vivo^18^.

Here we report the effects of SMQ-03-20 on age-related clustering of endogenous mPDE11A4 in ghost axons of the mouse brain. Across sexes, NIA C57BL6 mice show a robust age-related increase in mPDE11A4 ghost axons in ventral subiculum that reduces by ∼50% following oral dosing with SMQ-03-20 (effect of age x treatment: F(1,16)=8.75, P=0.0093; Post hoc vs. old vehicle: young vehicle P=0.0002, old drug P=0.0018). mPDE11A4 ghost axons also significantly increase with age in vCA1 (effect of age: F(1,16)=57.45, P<0.0001), dorsal subiculum (effect of age: F(1,16)=0.0142) and dorsal CA1 (Two Way ANOVA equal variance failed; Rank Sum Test for effect of age: T(10,10)=75.0, P=0.0254); however, SMQ-03-20 has no significant effects in these brain regions. Strikingly, amygdala of old vehicle-treated mice demonstrates a ∼4-fold increase in mPDE11A4 ghost axons relative to young mice, and this is completely reversed by treatment with SMQ-03-20 (Two Way ANOVA fails normality; ANOVA on Ranks for effect of group: H(3)=9.57, P=0.0226; Post hoc vs young vehicle: only old vehicle reached P<0.05).

**Figure 6.**
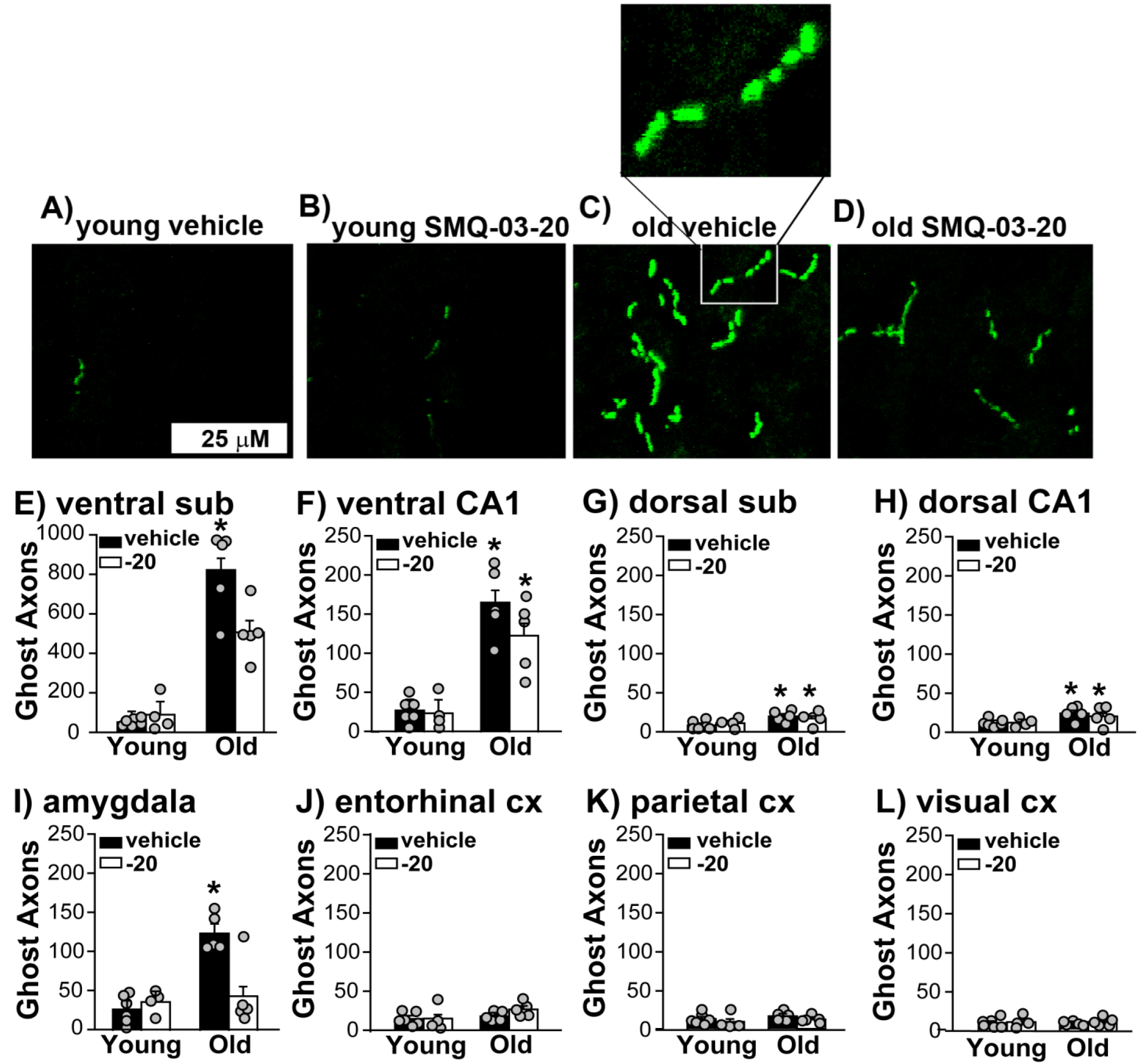
A single oral dose of 30 mg/kg SMQ-03-20 reverses age-related clustering of endogenous mPDE11A4 in the female and male mouse brain. Exemplar images of mPDE11A4 immunofluorescence from ventral subiculum (sub) of A) young vehicle-treated, B) young SMQ-03-20-treated, C) old vehicle-treated, and D) old SMQ-03-20-treated NIA C57BL6 mice (female shown). Note that upon higher magnification, many mPDE11A4 ghost axons reveal themselves to be a linear trail of spherical dots as opposed to a continuous filament (see C inset and ^1^). Quantification of mPDE11A4 ghost axons in mPDE11A4-expressing hippocampal subfields, including A) ventral subiculum (sub), B) ventral CA1, G) dorsal subiculum, and H) dorsal CA1. Quantification of mPDE11A4 ghost axons in brain regions that receive hippocampal projections, including the I) amygdala, J) entorhinal cortex (cx), K) parietal cortex, versus one that does not L) visual cortex. *vs young, P<0.05-<0.0001; #vs old vehicle, P=0.0018.

## DISCUSSION

Here, we provide evidence that age-related overexpression of PDE11A4 triggers LLPS (a.k.a. de-mixing or biomolecular condensation) of the enzyme and PDE11A4 small molecule inhibitors reverse this LLPS both *in vitro* and *in vivo*. Computational analyses reveal that mPDE11A4 and hPDE11A4 share many “pro-LLPS” sequence features, including—but not limited to—intrinsically disordered regions, non-covalent pi-pi interactions, prion-like domains, and a highly favorable charge patterning κ score (Figure 1). In HT22 and COS1 cells, we show that age-related mPDE11A4 spherical clusters are membraneless droplets that progressively fuse with each other over time in a concentration-dependent manner (Figure 2). Consistent with these *in vitro* observations, we see in the mouse brain that age-related clustering of mPDE11A4 in so-called “ghost axons” often reveals itself upon higher magnification to be a trail of adjacent spheres as opposed to a continuous filament-like structure (Figure 5; ^1^). Our PDE11A4 inhibitors then reverse this age-related LLPS of PDE11A4 (i.e., re-mix; Figure 3), with PDE11A4 able to de-mix once again after a 5-hour washout of less effective/lower concentrations of a PDE11Ai (Figure 4). Thus, PDE11A4 exhibits the main features of LLPS ^23, 25^ in that it forms membraneless spherical droplets that progressively fuse over time in a concentration-dependent and reversible manner.

### Computational analyses identify numerous LLPS-promoting features in both mPDE11A4 and hPDE11A4

In addition to its previously described N-terminal IDR and pS117/pS124 signature within that IDR that drives age-related clustering^1^, we show here that PDE11A4 exhibits many other biophysical/physiochemical properties that are associated with proteins that undergo LLPS^24, 25^. Like alpha-synuclein ^24^, CIDER^39^ characterizes PDE11A4 as a weak polyampholyte and weak polyelectrolyte that likely forms globules (i.e., category “R1” on the Das Pappu diagram; Figure S1). That said, the formation of PDE11A4 globules may be context dependent (i.e., category “R2”) given the fact that its FCR is 0.24223, which is just below the R2 category cutoff of 0.25. Notably, tau is an R2 LLPS protein ^24^. Such context dependence would be consistent with the fact that LLPS of PDE11A4 increases with age, perhaps in response to age-related drops in brain pH ^44^, and our catalytic inhibitors can reverse PDE11A4 LLPS (Figures 3-4). The suggestion of context-dependent condensation is also consistent with the PhaSePred^5^ prediction that PDE11A4 is likely to undergo both homotypic and heterotypic LLPS.

Experimental evidence to date aligns with the suggestion that PDE11A4 is capable of both homotypic and heterotypic LLPS under physiological conditions. Regarding homotypic condensation, post-translational modifications of IDRs are known regulators of homotypic LLPS ^25^, and the PDE11A4-pS117/pS124 signal that drives age-related clustering of PDE11A falls within its N-terminal IDR^1^. The fact that age-related LLPS occurs in parallel with more PDE11A4 protein being associated with the membrane fraction^1^ supports the possibility of PDE11A4 also undergoing heterotypic LLPS. Even though LLPS condensates themselves are membraneless, heterotypic LLPS often takes place at the plasma membrane in response to ing proteins being recruited there following receptor activation ^45, 46^. Consistent with this model, PDE11A4 is not directly inserted into the membrane but rather is indirectly associated with the membrane^1^. It will be of great interest to future studies to methodically determine the mechanistic underpinnings of PDE11A4 LLPS.

### PDE11A4 meets the minimum requirements for undergoing LLPS in cells

At a minimum, proteins that undergo LLPS should form membraneless spherical droplets that fuse over time in a concentration-dependent and reversible manner^23, 25^, and we show here PDE11A4 meets all of these criteria (Figure 2-4). PDE11A4 spherical droplets being membraneless is highly consistent with previous studies showing PDE11A4 clusters fail to colocalize with any membrane-bound organelle markers tested to date ^1^. No other PDEs have been reported to undergo LLPS to date, but that may not be surprising given that most do not increase in expression with age and many actually decrease in expression across the lifespan^1, 16, 18, 30, 34, 47, 48^. That said, the regulatory R1α subunit of protein kinase activated by cAMP (PKA) undergo LLPS in response to elevated cAMP levels^49^. The fact that LLPS appears to be responsible for age-related clustering of PDE11A4 is highly interesting because LLPS is gaining interest in the mechanistic study of dementia and neurodegenerative diseases ^22, 23, 44^ ^50–52^. In this context, it may be interesting to note that 2 rare PDE11A variants have been associated with early-onset Alzheimer’s disease^53^ and exacerbated age-related increases in PDE11A expression have been associated with dementia in elderly traumatic brain injury patients^1^. Interestingly, many factors that regulate LLPS in general have been reported to change in the aging and AD brain ^1, 23, 24, 44, 54–57^. These factors include elevated protein concentrations as well as reduced salt concentration, temperature, and pH ^1, 44, 54–57^. A possible relationship between aging/AD-related decreases in brain pH and PDE11A4 LLPS are particularly plausible given the established interplay between cAMP/cGMP/PDE signaling and pH-regulated processes (e.g., ^58–62^).

### PDE11Ai’s inhibit mPDE11A4 catalytic activity and reverse mPDE11A LLPS

Here we showed that 5 PDE11Ai’s across 3 different scaffolds robustly reversed mPDE11A4 LLPS (i.e., re-mixed the enzyme). For the most part, a 5-hour washout of low, but not high, concentrations of the compounds led to mPDE11A4 de-mixing once again. This raises the interesting possibility that there may be a high versus low affinity binding site for these PDE11Ai’s, with binding to the high affinity site being more readily reversible and the low affinity site not being reversible. There is precedent in the PDE4 family, with some isoforms including both a high affinity and low affinity binding site for inhibitors^63^. In this regard, it is interesting to note that PDE11A4 binds cGMP both as a substrate at the catalytic site and at the regulatory GAF-A domain, the latter of which also binds inhibitors of PKG (c.f., ^64^). We believe this is particularly intriguing given that catGRANULE predicted the GAF-A domain to have LLPS propensity (Figure 1).

As previously reported in C57BL/6J and BALB/cJ mice^1^, female and male NIA C57BL6 mice show a robust age-related increase in mPDE11A4 ghost axons in ventral subiculum and vCA1. In contrast to this previous report, NIA C57BL6 mice also exhibited age-related increases in mPDE11A4 ghost axons in dorsal subiculum and dorsal CA1. Interestingly, only the age-related increase in ventral subiculum was reduced by the single oral dose of SMQ-03-20 (∼50% reduction). We also examined brain regions that receive hippocampal projections and found that the amygdala, but not the entorhinal or parietal cortex, shows a robust age-related increase in mPDE11A4 ghost axons that is completely reversed by the SMQ-03-20 treatment (Figure 5). This brain region-selective effect of the inhibitors *in vivo* is striking and raises the possibility that the effectiveness of PDE11Ai’s to reverse LLPS may be modulated by the presence of another binding partner that is expressed in some but not all of these regions (i.e., homotypic versus heterotypic condensation).

The mechanism by which PDE11Ai’s re-mix PDE11A4 remains to be determined. It is not by reducing PDE11A4 protein expression, as we found no effects of the PDE11Ai’s on PDE11A4 protein expression (Table 2). It may be possible that PDE11Ai’s re-mixing PDE11A4 droplets by virtue of having elevated cAMP/cGMP levels (i.e., by inhibiting PDE11A4 catalytic activity). That said, increased cAMP promotes LLPS of the PKA R1α subunit^49^ and select PDE4 inhibitors trigger PDE4A4 to adopt an LLPS-like phenotype (i.e., it accumulates in membraneless spherical structures)^65–67^. It is possible that PDE11Ai’s ultimately decrease PDE11A4-p117/p124 (that is the post-translational modification that promotes PDE11A4 LLPS^1^). That said, preventing phosphorylation of S117/S124 by introducing an alanine mutation at each site only partially reduces PDE11A4 LLPS, it by no means eliminates it as do higher concentrations of the PDE11Ai’s (Figures 3-4). Another intriguing possibility is that application of the PDE11Ai’s reduces PDE11A4 packaging/repackaging via the trans-Golgi network since we previously showed that stimulating Golgi packaging increased PDE11A4 LLPS^36^. Indeed We are actively investigating these possibilities and more.

### Conclusion

Depending on the molecule, LLPS can sequester unneeded protein, buffer proteins (i.e., temporarily store and then release upon demand), or accelerate biochemical reactions by virtue of concentrating enzymes with substrates in membraneless organelles ^23^. It will be of great interest to determine what function PDE11A4 LLPS is serving in the aging brain so that we may fully understand how PDE11A4 should be targeted for therapeutic gain. We recently reported that a biologic that disrupts homodimerization of mPDE11A4 (i.e., an isolated PDE11A4 GAF-B domain) reversed age-related clustering of PDE11A4 protein both in vitro and in the old mouse brain and rescued age-related social memory deficits ^15, 26^. This biologic also selectively degraded membrane-associated mPDE11A4, thereby increasing the mPDE11A4 cytosolic:membrane ratio as we saw here with PDE11Ai’s. This may suggest that the ability of our PDE11Ai’s to reverse PDE11A4 LLPS and increase the mPDE11A4 cytosolic:membrane ratio will add to the therapeutic effects of their enzymatic inhibition.

## ACKNOWLEDGEMENTS

The authors would like to thank Latarsha Porcher and William Capell for technical assistance with the live imaging study, as well as Asmaa Hijazi for assistance with making peanut butter pellets. The authors would also like to acknowledge Jeremy Eberhard, Dennis Colussi, John Gordon, and Wayne Childers for generating in vitro screening data reported elsewhere ^18^ that contributed to decisions of which compounds we tested herein.

